# PD1 is transcriptionally regulated by LEF1 in mature T cells

**DOI:** 10.1101/2023.03.01.530714

**Authors:** Pin Zhao, Lanming Sun, Cong Zhao, Samiullah Malik

**Affiliations:** Department of Clinical Laboratory, The Third People’s Hospital of Shenzhen, Southern University of Science and Technology, National Clinical Research Center for Infectious Diseases, Shenzhen, China; Department of Prevention, Health care and Fertility, Xinfuli Community Hospital, Linhongnong Road, Dahongmen, Fengtai District, Beijing, China; Department of Pathogen Biology, Shenzhen University Health Science Center, Shenzhen 518055, China

**Keywords:** PD1, LEF1, CD8+ T cells, Treg cells, Concanavalin A

## Abstract

Programmed cell death 1 (PD1) plays a vital role in cancer immune evasion. So monoclonal antibodies targeting PD-1 to boost the immune system are being developed for the treatment of cancer. But the effect of PD1/PD-L1 blocking antibody only works in a subset of patients or some tumors. It is well established that PD-1 can affect the positive selection of T cells. Mice lacking PD1 or PD-L1 had more DP T cells in the thymus. However, how it affects the T-cell repertoire is not very clear. For T cell development and maturation, TCF1 and its homolog LEF1, act as indispensable factors and TCF1 has been most studied. In our work, we found that LEF1, not TCF1, positively regulated the transcription of PD1 by generating two mice models, including ERT2-CRE-LEF1^flox/flox^ mice and concanavalin A (ConA)-mediated T activation mice model. We investigated LEF1 and PD1 expression in mature T cells, including CD4+ T cells, CD8+ T cells, and Treg cells in these two mice models. And we also evaluated the expression of PD1 in tumor-infiltrating lymphocytes in the tumor-implantation experiments. Furthermore, the regulation of PD1 by LEF1 was verified in LEF1 knockdown cells. Our conclusions provided a novel insight into PD1 regulation in immune responses and explored potential strategies for clinical anti-PD1 treatment.

## Introduction

Mature T cells originate from the thymus and are distributed to the peripheral lymphoid organs via the bloodstream, including the spleen, lymph nodes, pharyngeal tonsils, appendix, and lymphatic nodules and tissues, which are all over the body. These peripheral mature T cells are heterogeneous and roughly divided into three types, including naïve, effect, and memory T cells. Although the renewal of mature T cells in adults has decreased with age, the number of mature peripheral T cells remains constant from puberty and the lifespan of mature T cells is relatively long [1]. Mature T cells are essential for protecting the body against infections by pathogenic bacteria and viruses and cancerous cells, which involve a series of immune responses. In general, five main types of T cells with different functions are responsible for the T cell-mediated immune response, including cytotoxic T cells, helper T cells, regulatory T cells (Treg cells), natural killer T cells (NKT cells), and memory T cells. Cytotoxic T cells, also known as CD8+ T cells, can directly destroy cells infected by pathogens or cancer cells by making use of granules to induce targeted cell apoptosis. Helper T cells are a kind of CD4+ T cells. Once activation, they are differentiated into various effector subtypes, such as Th1, Th2, Th9, Th17, and Tfh, which secrete distinct cytokines to activate CD8+ T cells and B lymphocytes [2]. Natural killer T cells and regulatory T cells are a small part of CD4+ T cells. There are two types of regulatory T cells, natural Treg cells, and induced Treg cells. Both of them maintain a balance and function to suppress the immune response. In normal conditions, Treg cells play a crucial role in maintaining the potential auto-reaction of peripheral T cells [3]. NKT cells are a unique subpopulation of mature T cells that express both NK cell receptors (NK1.1) and incomplete T cell receptor (TCR) repertoire on the cell surface. By producing various cytokines, such as IL4 and IFNγ, NKT cells perform cytotoxic functions, the same as NK cells, in which there is the involvement in T helper types 1 and 2 (Th1 and Th2) responses [4-6]. Memory T cells are long-lived and antigen-specific T cells, including CD4+ T cells and CD8+ T cells. When it comes to intracellular pathogens or inflammation, memory T cells are quickly converted and exert effector functions in response to reencountered antigens. So, the population of memory T cells is a crucial factor in assessing the efficacy of vaccination [7].

The transcription factor T cell factor 1 (TCF1) and its homolog LEF1 are widely expressed in T cells and are indispensable for early T cell development [8]. Structurally, both TCF1 and LEF1 contain a histone deacetylase domain (HDAC) and a high mobility group (HMG) DNA-binding domain in the C terminus with all isoforms. While only the long isoforms of TCF1 and LEF1 contain a β-catenin-binding domain (β-BD) at their N-terminus, which suggests that TCF1 and LEF1 transcriptionally regulate downstream genes of the Wnt signaling pathway via the interaction with β-catenin. In the development of T cell maturity and T lineage commitment, the functions of TCF1 are relatively more versatile than those of LEF1, perhaps because TCF1 transcriptionally represses the expression of LEF1. It has been evidenced that TCF1 promotes Th2 and inhibits Th1 and Th17 differentiation through the Wnt signaling pathway [9,10]. However, the roles of TCF1 and LEF1 in mature T cells have been considered to be beyond their interactions with β-catenin and Wnt signaling pathway. For example, it has been reported that the interaction between TCF1/LEF1 and Runx3 is required for silencing cd4 expression in CD8+ T cells [11]. LEF1 interacts with GATA3 in Th2 cells [12]. And TCF1 binds Foxp3 to repress IL2 expression in Treg cells [13]. Recent research has shown that TCF1 and LEF1 positively regulate the immune-suppressive functions of Treg cells by inducing the expression of the signature genes, such as Ikzf4 and Izumo1r [14].

Programmed cell death 1 (PD1) is a transmembrane protein and abundantly expressed in various effector immune cells, especially activated T cells. PD1, known as an immune checkpoint, negatively regulates the immune response. Specifically, PD1 is composed of an immunoglobulin variable region-like domain in its N-terminus, a transmembrane domain, and three tyrosine-based signal motifs (two SHP1 and one SHP2) in the C-terminus. The recruitment of its receptor programmed cell death 1 ligand 1 (PDL1) promotes T cell inactivation, which has been clinically interfered with within cancer immunotherapy. In terms of molecular mechanisms, after the TCR signal is activated by antigen recognition, the motifs of PD1 are phosphorylated by LCK [15]. The expression of PDL1 is dynamically upregulated in cancer cells. When PD1 binds to PDL1, these motifs are dephosphorylated, resulting in terminating TCR signal transduction and T cell activation and thus promoting cancer cells to escape from the immune response in the tumor microenvironment [16]. In addition, PD1 can repress T cell activation in an indirect manner. CK2 increases and phosphorylates PIP3 in activated T cells [17]. It has been reported that the engagement of PD1 downregulates CK2 expression and induces PIP3 dephosphorylation for PI3K inactivation [18]. Meanwhile, PD1 inhibits T cell proliferation by decreasing the expression of CDKs and the ubiquitin ligase, SCF^skp2^ [19,20]. Therefore, PD1, as well as LAG3, CD39, CTLA4, and TIGIT, are considered T-cell exhaustion markers. CD8+ T cells with the expression of PD1 impair the antiviral motility and effect functions, while they can also prevent the overactivation of antigen-specific CD8+ T cells [21]. However, PD1 is highly expressed in Treg cells originating from activated T cells [22]. And its expression improves the suppressive capacity of Treg cells.

In the present study, we found that LEF1 transcriptionally regulated PD1 expression in mature T cells, such as CD8+ T cells and Treg cells. LEF1 knockout repressed PD1 expression and thus influences the immune-suppressive capability of Treg cells. By contrast, ConA-mediated T cell activation induced LEF1 expression, and thus it upregulated PD1 expression in CD8+ T cells and Treg cells. It provided a novel insight into PD1 regulation in immune responses and immunotherapy.

## Materials and Methods

### Animal experiments

C57BL/6 mice, ERT2-CRE transgenic mice, and LEF1^flox/flox^ mice were purchased from the Jackson Laboratory. Generation of ERT2-CRE-LEF1^flox/flox^ mice was performed by backcrossing ERT2-CRE transgenic mice and LEF1^flox/flox^ mice for >6 generations. All mouse experiments were carried out with almost equal numbers of mice at same age (4-6 weeks), and no disability or defects was done. All experiments were approved by the local and federal laws on animal research.

For induction of ERT2-CRE-mediated recombination, mice were settled by intraperitoneal injection of Tamoxifen (Sigma-Aldrich, #T5648, dissolved in corn oil and at a concentration of 20 mg/ml) for 5 consecutive days, using approximately 75 mg/kg body weight. C57BL/6 mice were injected with 50 ul concanavalin A (Con A) at a concentration of 5 μg/ml for four consecutive days.

### In vitro cell stimulation

Single cell suspension from the mouse spleen was isolated and cultured in RPMI 1640 medium (SH3080902, Cytiva) plus 10% fetal bovine serum (A3160801, Gibco) and 50 μg/ml penicillin and streptomycin (100 units/ml, SV30010, HyClone). 1× 10^6^ cells were counted and treated with concanavalin A at a final concentration of 5 ng/ml. Cells were cultured for 7 days in a 5% CO2 incubator at 37°C and detected with anti-CD25 (eBiosciences, PC61.5). The stained cells were analyzed by the CytExpert Platform (Beckman).

### Cell culture

Human MOLT4 cells (CC1908, cellcook) were cultured in RIPM (SH3080902, Cytiva), 10% fetal bovine serum (A3160801, Gibco), penicillin (100 U/ml) /streptomycin (100U/ml) (SV30010, HyClone).

### LEF1 knockdown

LEF1-specific targeted sequences were chosen according to online shRNA design tools of Invitrogen http://www.invitrogen.com/rnai using the reference sequences of LEF1 (Gene Bank Accession No. AF288571.1).LEF1(LEF1-A : 5‘-CCGGCCATCAGATGTCAACTCCAAATTCAAGAGATTTGGAGTTGACATCTGATGGTT TTT-3’ and LEF1-B: 5’-CCGGGCTGGTCTGCAAGAGACAATTTTCAAGAGAAATTGTCTCTTGCAGACCAGCTTTTT-3’) were chemically synthesized and constructed into lentivirus vector (pLKO.1-GFP-ShRNA), respectively. The invalid RNAi sequence (5’-CCGGCAACAAGATGAAGAGCACCAATTCAAGAGATTGGTGCTCTTCATCTTGTTTTT-3’) was used as negative control. The sequences of plasmids with siRNAs were further confirmed by sequencing. The shRNAs for the first and the second exon of Lef1, were purchased from Sangon Biotech Co. Ltd (Shanghai, China). The shRNAs were cloned separately into LentiCRISPRv2-GFP (modified from Addgene) and transiently co-transfected with psPAX2 (12260, Addgene)and pMD2.G(12259, Addgene)in MOLT4 cells for 24 hours. GFP-positive cells were selected by fluorescence-activated cells sorting (FACS) flow cytometer (BD Biosciences). Cells were then cultured and reselected for stable GFP-positive cells. The expression of LEF1 were analyzed by Western blot and RT-qPCR.

### Western blot

Cells were harvested and washed twice with PBS. And then cells were suspended in 400 μl RIPA buffer (150 mM NaCl, 1% Nonidet P40, 0.5% sodium deoxycholate, 0.1% sodium dodecyl sulphate, 20 mM Tris-HCl, pH 7.6) containing 1 mM PMSF (SERVA) and protease inhibitory cocktail (Roche). Cell lysis was performed on ice for 25 min and then precleared by centrifugation at 14000 g for 10 minutes at 4°C. The supernatants were mixed with 6 x Laemmli buffer (40% glycerol, 240 mM Tris/HCl, pH 6.8, 8% SDS, 0.04% bromophenol blue, 5% β-mercaptoethanol) and denatured at 96ºC for 10 minutes. For nuclear protein, the harvested cells were incubated on ice for 30 minutes in 400 μl lysis buffer (20 mM Tris-HCl, pH 7.5, 0.5 mM EDTA, 150m M NaCl, 0.5% Nonidet P40, and 1 mM PMSF and protease inhibitor cocktail). After centrifugation at 14000 g for 10 minutes at 4°C, the pellets were collected and dissolved in 6 x Laemmli buffer. The boiled proteins were resolved by SDS-PAGE and transferred to nitrocellulose membranes (Millipore). The membranes were blocked in TBST (Tris-buffered saline, 0.075% Tween 20) with 5% fat-free milk for 1 h at room temperature and then incubated with primary antibody overnight at 4°C. After washing with TBST three times, the membranes were incubated with secondary antibody for 1 hour at room temperature. Bound secondary antibody conjugated to horseradish peroxidase (HRP) is visualized for chemiluminescent detection according to an ECL detection kit (Beyotime Biotechnology). The primary antibodies used were: TCF1 (Dilution of 1:5000, C64C7, Cell Signaling); LEF1 (Dilution of 1:5000, C12A5, Cell Signaling); β-actin (Dilution of 1:2000, E0919, Santa Cruz); PD1 (Dilution of 1:2500, 66220-1-Ig, Proteintech).

### Luciferase report assay

Different groups of 293T cell were cultured in 24-well plates in triplicate. After 24 h, 293T cells were transfected with PGL3 firefly luciferase reporter vector (Promega) containing wild-type and its mutant of PD1 promoter region (WT and Mut), pCMV beta mammalian expression vector containing full-length human LEF1, and pRL-TK Renilla plasmid using lipofectamine 3000 reagent (Thermo Fisher, USA, No.L3000015). The concentrations of PD1 promoter region and control vector were optimized at same level before the transfection into HEK293T cells. Luciferase and Renilla signals were measured 48 h after transfection by a Dual-Luciferase Reporter Assay Kit (Promega, No. E1980). Data were analyzed and normalized by the division of firefly luciferase activity with that of Renilla luciferase to eliminate transfection efficiency difference.

### Histochemical staining

For the hematoxylin and eosin (H&E) staining, after being deparaffinized and hydrated, paraffin-embedded spleen tissue sections were stained with H&E for histopathological examination. The clinical scoring with H&E staining was adapted from a modified Ishak system.

For immunofluorescence staining, paraffin-embedded spleen tissue sections were deparaffinized and hydrated before processed by antigen thermal retrieval. The sections were then incubated with primary antibody (anti-mouse PD1, 66220-1-Ig, Proteintech) at 4°C overnight followed by being infiltrated with DAPI (C1005, Beyotime Biotechnology) to stain the cellular nuclei.

### RNA extraction and RT-qPCR

RNA was extracted and purified with Total RNA purification (Ezymo) according to the manufacturer’ instructions. 1 μg RNA was used for cDNA with SuperScript III Reverse Transcriptase kits (Life Technologies). The Real-time quantitative PCR reactions were performed by which cDNA with a dilution of 50 or 100-fold, was mixed with a DNA SYBR Green I Master Mix (Roche) on a LightCycler machine (Roche). All gene expression levels were normalized to GAPDH. Primer sequences were presented in **Table 1**. The experiments for statistical analysis were performed in triplicates and determined by two-tailed Student’s t-test.

**Table 1.**
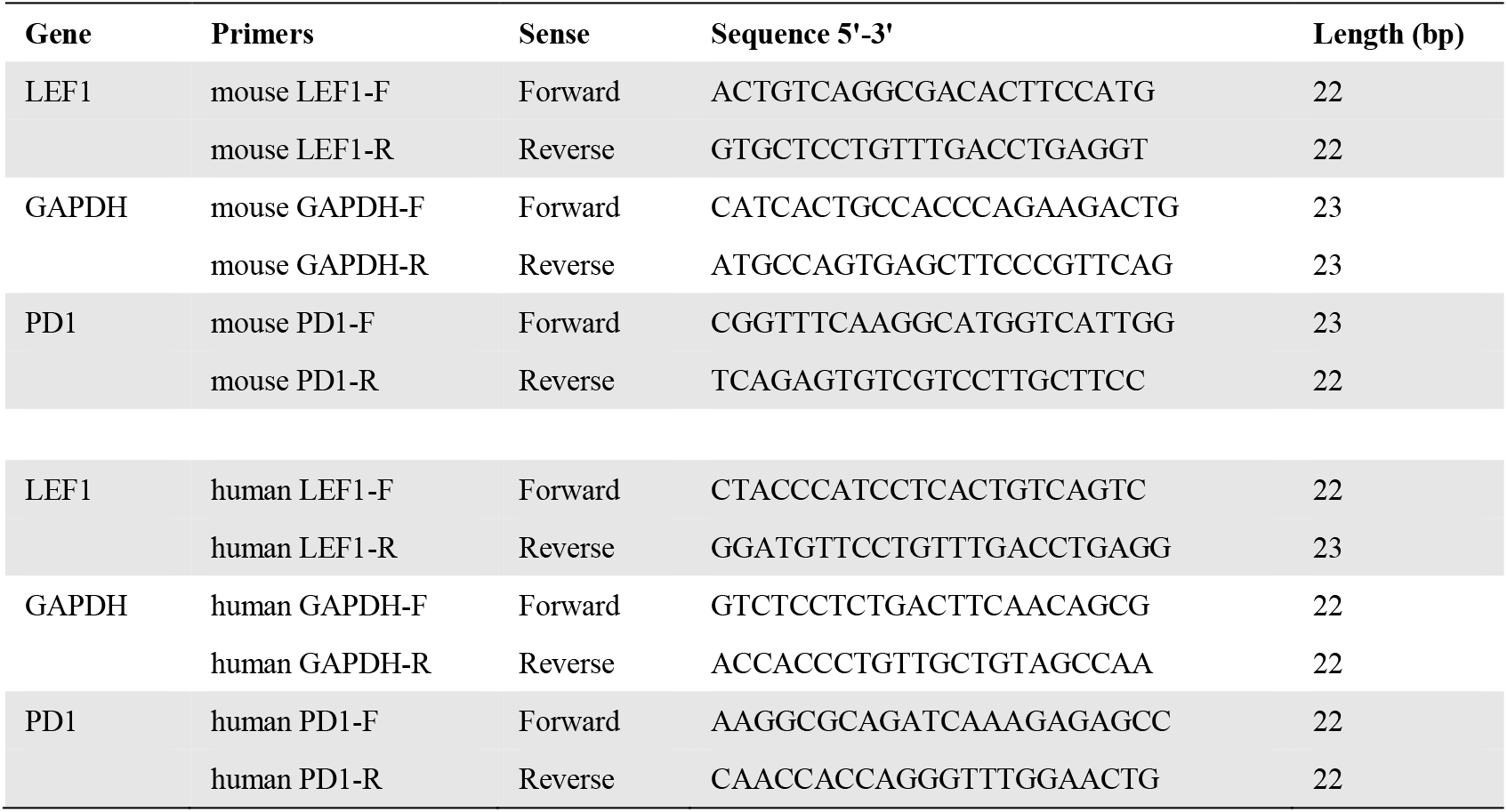
The list of qPCR primers.

### Flow cytometry and cell sorting

Cells were prepared from the mouse thymus, spleen, and B16F10 tumors and surface-stained with different combinations of all fluorescent-labeled antibodies from eBiosciences: anti-TCRβ (H57–597), anti-CD4 (M1/69), anti-CD8α (53-6.7), anti-CD45.1 (A20), anti-CD44 (IM7), anti-CD25 (PC61.5). For intranuclear staining, cells were firstly stained by surface antibodies and then fixed and permeabilized with Foxp3/transcription factor staining buffer set (eBiosciences), following anti-Foxp3 (eBiosciences, FJK-16s) staining. The stained cells were analyzed by the CytExpert Platform (Beckman). All the antibodies were used mouse cells with a dilution of 1:50. The results were directly analyzed by CytExpert Software (Beckman).

### ChIP and ChIP-qPCR

The ChIP assay was performed following the manufacturer’s instructions (Abcam). In brief, mouse embryonic stem cells were fixed with 0.75% formaldehyde directly to cell culture media at room temperature for 10 min, followed by incubating with 125 mM glycine for 5 min at room temperature. Cells were washed with cold PBS, scraped and resuspended in lysis buffer (50 mM HEPES-KOH pH7.5, 140 mM NaCl, 1 mM EDTA pH8, 1% Triton X-100, 0.1% Sodium Deoxycholate, 0.1% SDS) with protease inhibitor cocktail (SERVA). Chromatin was then sonicated to the fragments of 0.5 kb and preincubated with the Dynabead G (Merck Millipore) and anti-LEF1 antibody overnight at 4°C. The beads without antibody were as control of IgG. The chromatin bound to the beads was eluted in 500 μl of freshly prepared elution buffer (1% SDS, 0.1 M NaHCO3). After reversing the cross-linking, the samples were deproteinized and phenol–chloroform-extracted, and DNA was ethanol-precipitated using glycogen as a carrier. Pellets were resuspended in 100 μl of H2O for qPCR analysis.

### Statistical analysis

The data were presented as mean ± SD value. Differences in the results of two groups were evaluated using either two-tailed Student’s t-test or one-way ANOVA. The results with p < 0.05 were considered statistically significant. Statistical analyses were performed with the software GraphPad Prism 8.0 (GraphPad Software, San Diego, CA, USA) was used to plot graphs.

## Results

### PD1 expression was reduced in LEF1 ^-/-^ mice

In early stages of thymocyte maturation, TCF1 is induced by Notch signaling and it stimulates β-selection for CD4+CD8+ double positive stage [23-25]. In order to become single CD4+ or CD8+ T cells, TCF1 and LEF1 cooperatively suppress CD4+-lineage through positive regulation of the transcription factor Th-POK and interaction with RUNX3 [11]. It has been reported that TCF1^−/–^LEF1^−/–^ CD8+ T cells display a decreased expression of CD4+ signature genes, such as *Cd40lg, St8sia6, Lgmn* and *Itgb3* [26]. Although ablation of TCF1 and LEF1 cannot alter the distribution of memory and effect subsets of Treg cells, TCF1 and LEF1 are essential for maintaining immunosuppressive function of Treg cells [14].

To define intrinsic regulation of LEF1 in T cells, we generated mice with a genetically-engineered mouse model for global, inducible deletion of LEF1 by crossing LEF1^flox/flox^ to ERT2-Cre mice. In this model, ERT2-Cre is conditionally activated by tamoxifen (TA) administration (**Figure 1A**). And 250 ug tamoxifen was administered intragastrically every day for 5 days to induce whole-length LEF1 deletion. After 5 days of tamoxifen treatment, we isolate spleens and lymph nodes for further analysis. Compared with mice without tamoxifen treatment, LEF1 expression was decreased in both of spleen and lymph nodes by western blot (**Figure 1B**). It is suggested that efficient Cre expression and LEF1 deletion in ERT2^Cre^ LEF1^flox/flox^ mice. Meanwhile, PD1 expression was also reduced, which was consistent with LEF1 expression. Furthermore, we observed that LEF1 deletion scarcely had an effect on the morphology and composition of mice spleen, while PD1 expression was lower than that in LEF1 wildtype mice by immunohistochemistry (**Figure 1C**). These results indicate LEF1 ablation impacts PD1 expression in mice.

**Figure 1.**
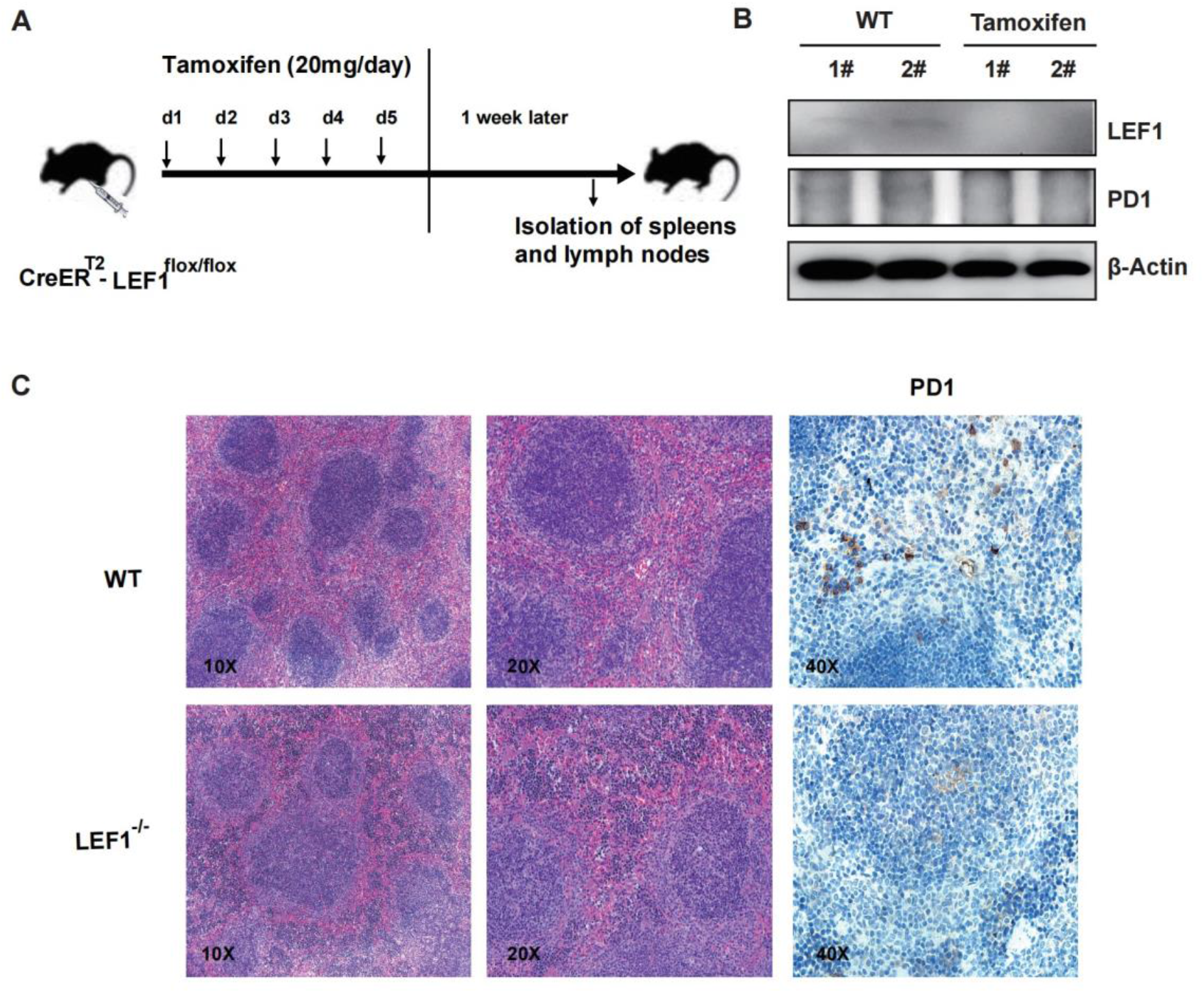
LEF1 knockout reduced PD1 expression. (A) The mice model for LEF1 knockout. The ERT2-Cre-LEF1^flox/flox^ mice were administered intragastrically with 250 ug tamoxifen for five days and continued to culture the mice for another week for further analysis. (B) Western blot of LEF1 and PD1 in mice spleen and lymph nodes treated with or without tamoxifen. (C) Hematoxylin and eosin (HE) staining and immunohistochemical staining of PD1 in mice spleens with different magnifications.

### PD1 in CD8+ T cells and Treg cells were downregulated in LEF1^-/-^ mice

LEF1 is mainly expressed in all kinds of T cells with different levels, while PD1 is found to be highly expressed in exhausted CD8+ T cells, and also in activated T cells, such as Treg cells [27,28]. So, we checked PD1 levels in CD8+ T cells and Treg cells and found that PD1 decreased in the CD8+ T cells from spleens and lymph nodes of ERT2-Cre-LEF1^flox/flox^ mice compare of that in wild-type mice (**Figure 2A and 2B**). And Treg cells in ERT2-Cre-LEF1^flox/flox^ mice also displayed a lower level of PD1 expression than that in wild-type mice (**Figure 2C**). It suggested that PD1 expression was influenced by LEF1.

**Figure 2.**
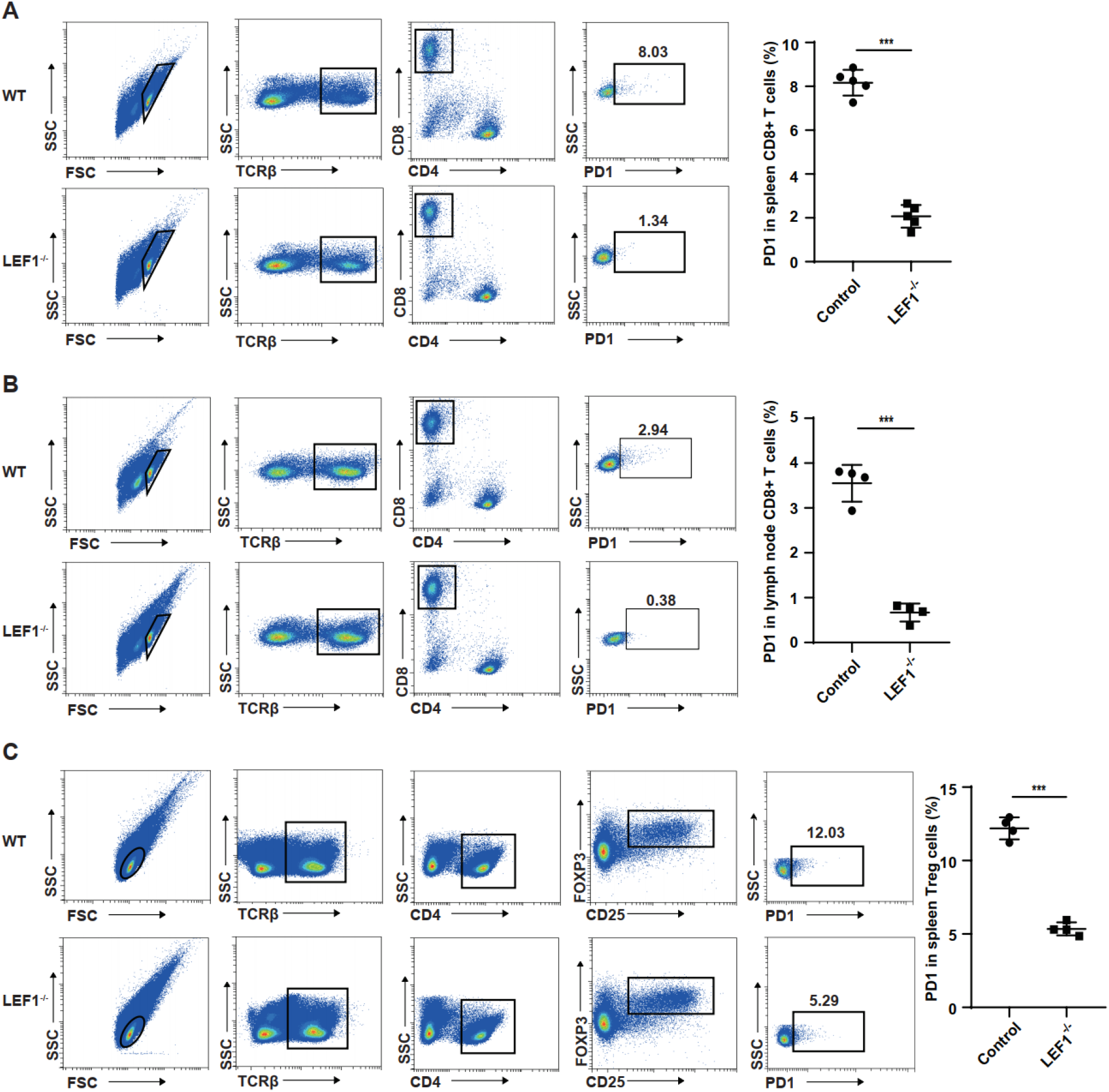
PD1 was reduced in different types of T cells. (A) Flow cytometry analysis of TCRβ+ CD8+ PD1+ T cells in the spleens of wild-type mice and LEF1 knockout mice. Bars in scatter plots indicate median, error bars indicate interquartile range. (B) Flow cytometry analysis of TCRβ+ CD8+ PD1+ T cells in the lymph nodes of wild-type and LEF1 knockout mice. (C) Flow cytometry analysis of TCRβ+ CD4+ CD25+ FOXP3+ PD1+ T cells in the spleens of wild-type and LEF1 knockout mice. TCRβ+ CD4+ CD25+ FOXP3+ T cells represent Treg cells. The data were shown as mean ± SD of triplicates experiments. *p < 0.05, **p < 0.01, *** p < 0.005 compared with the corresponding control. Differences in the results of two groups were evaluated using either two-tailed Student’s t-test or one-way ANOVA.

### ConA-stimulated T activation induced the expressions of LEF1 and PD1

To further verify whether LEF1 regulates PD1 expression, we used another mice model to verify it. Mice were treated with concanavalin A (ConA) with a low dose for four times every other day, which induced T cell activation (**Figure 3A**). Concanavalin A is firstly discovered from jack bean Canavalia ensiformis and extensively binds to sugar residues on the cell surface of various cell types [29,30]. High doses of ConA is usually used to establish the mice model of acute hepatocyte damage by intravenous injection, which induced abundant inflammatory cytokines. While low dose of ConA can stimulate immune activation and memory [31,32]. It has been reported that ConA, as well as phytohemagglutinin (PHA), activates T cells by binding to the T cell receptor (TCR)-CD3 complex [33]. And ConA has evidenced to extensively stimulate T cell proliferation in vitro [34]. By western blot, we found that after ConA stimulation, LEF1 expression is upregulated compared to wild-type mice (**Figure 3B**). To further verify it, we isolated T cells from spleens of wildtype mice and incubated with ConA for days ranging from 1 to 6, respectively. Flow cytometric analysis of these cells revealed that the frequency of CD25 positive T cells were gradually increased, along with the upregulated expressions of LEF1 and PD1 by RT-qPCR (**Figure 3C and 3D**). In mice, we detected PD1 expression in CD8+ and Treg cells followed by ConA through intraperitoneal injection. It showed that both PD1 in these two kinds of T cells were distinctly augmented (**Figure 3E and 3F**). Taken together, we concluded that LEF1 positively regulated PD1 expression in CD8+ and Treg cells.

**Figure 3.**
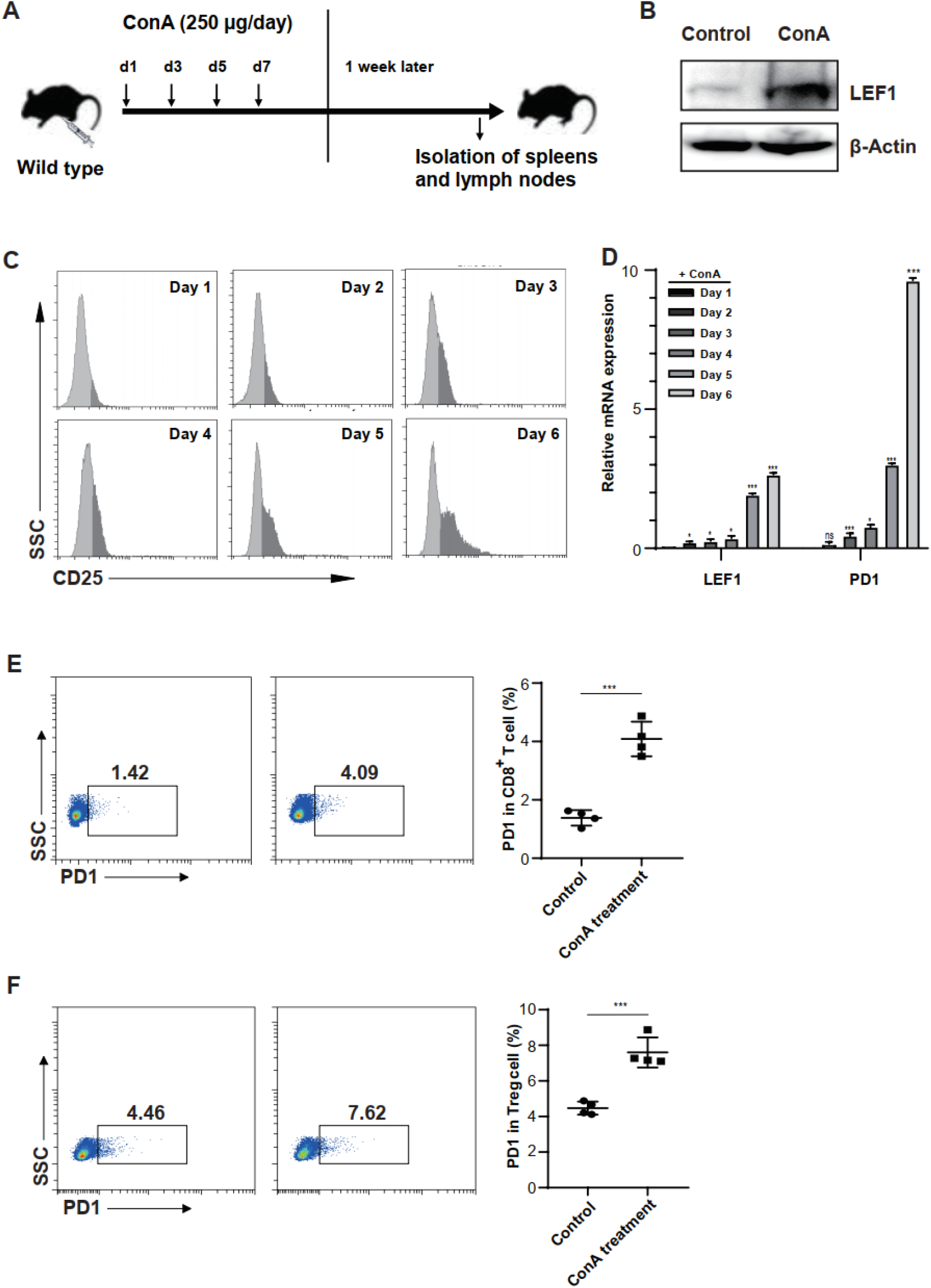
ConA stimulated the expressions of LEF1 and PD1 in T cells. (A) The mouse model stimulated by concanavalin A (ConA). (B) Western blot of LEF1 in mice spleens with or without ConA stimulation. (C) In vitro stimulation of T cells by ConA. CD25+ activated T cells were traced for 6 days and the mRNA levels of LEF1 and PD1 were measured by RT-qPCR (D). (E) Flow cytometry analysis of TCRβ+ CD8+ PD1+ T cells in the spleens of wild-type and ConA-stimulated mice. Bars in scatter plots indicate median, error bars indicate interquartile range. (F) Flow cytometry analysis of TCRβ+ CD4+ CD25+ FOXP3+ PD1+ T cells in the spleens of wild-type and ConA-stimulated mice. The data were shown as mean ± SD of triplicates experiments. *p < 0.05,**p < 0.01, *** p < 0.005 compared with the corresponding control. Differences in the results of two groups were evaluated using either two-tailed Student’s t-test or one-way ANOVA.

### PD1 was upregulated in ConA-stimulated tumor infiltrating lymphocytes

With clinical drug development for blocking the PD1 immune checkpoint in cancer immunotherapy because the interaction between PD1 and its receptor PDL1 induces immunological turnoff, it is also important to know whether PD1 is regulated by LEF1 in tumors. We implanted melanoma cells (B16F10 cells) into wild-type, ERT2-Cre-LEF1^flox/flox^ mice and ConA-treated mice. These mice have been treated with corn oil, tamoxifen, and ConA treatment before tumor implantation, respectively. Our analysis of FACS displayed that the ratio of CD4+ and CD8+ T cells was diverse among these three experiment groups (**Figure 4A**). Due that LEF1 is essential for T cell development, there was no tumor-infiltrating lymphocytes (TILs) in LEF1 deficiency mice, such as CD4+ and CD8+ T cells. However, the frequency of CD4+ T cells notably was higher in ConA-treated mice that that of wild-type mice. There was no statistical difference in CD8+ T cells between ConA-treated mice and wildtype mice. It suggested that it is unlikely to detect PD1 expression in ERT2-Cre-LEF1^flox/flox^ mice-implanted tumors. Therefore, we nearly cannot detect PD1 expression in such few CD8+ T cells from ERT2-Cre-LEF1^flox/flox^ mice (**Figure 4A**). But more PD1 was expressed in CD8+ T cells from ConA-stimulated mice than that from wild-type mice. Furthermore, PD1 was also detected in Treg cells. Our results showed that ConA stimulated PD1 expression, reaching 48.78%, almost twice that (29.25%) of the control mice (**Figure 4B**). Based on the results of these tumor models, we further validated that PD1 is regulated by LEF1 in CD8+ T cells and Treg cells.

**Figure 4.**
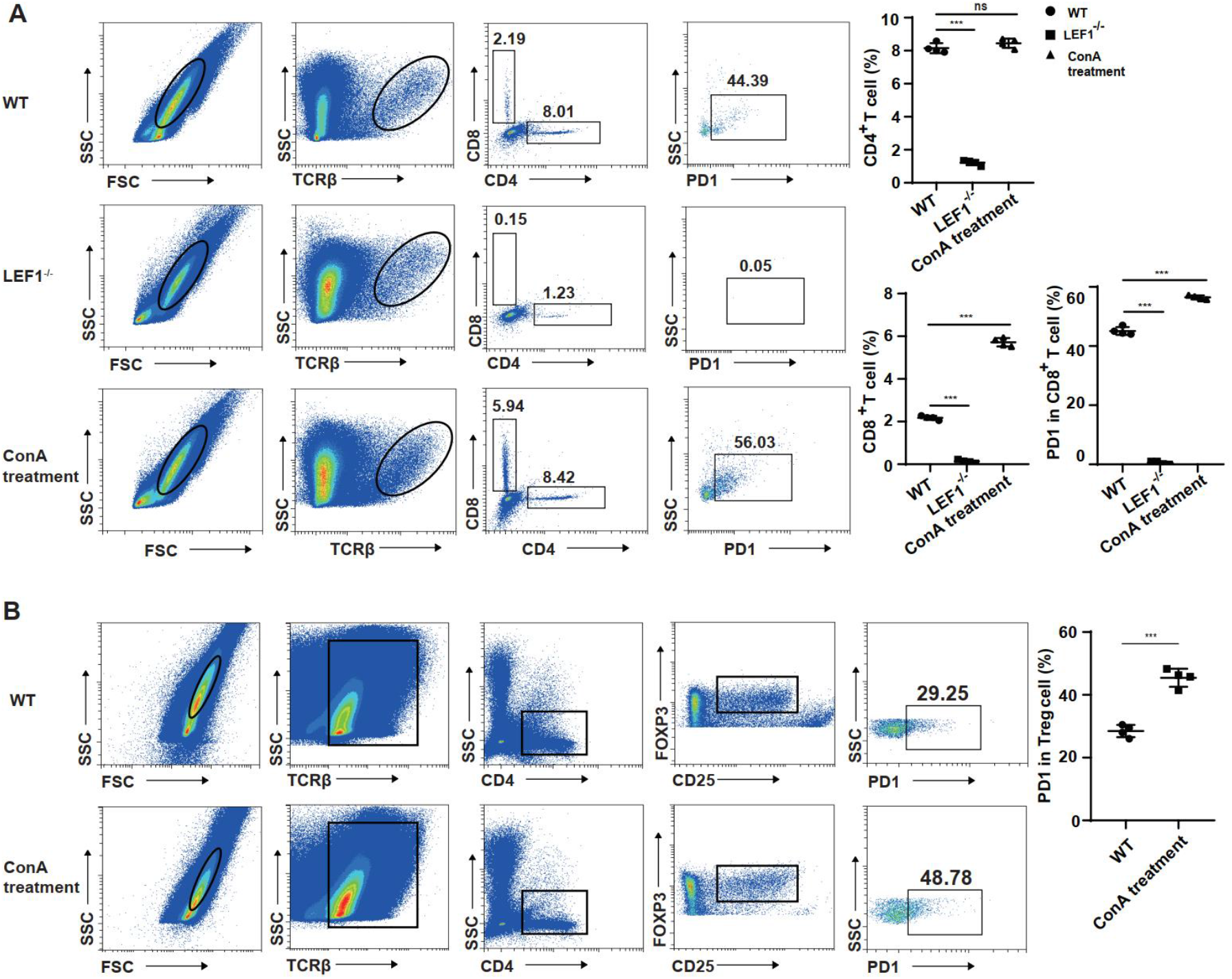
PD1 expression in tumor infiltrating lymphocytes. (A) Flow cytometry analysis of TCRβ+ CD4+, TCRβ+ CD8+ T cells and the frequency rate of PD1 in TCRβ+ CD8+ T cells in tumors of wild-type, LEF1 knockout and ConA-stimulated mice. (B) Flow cytometry analysis of TCRβ+ CD4+ CD25+ FOXP3+ PD1+ T cells in the tumors of wild-type and ConA-stimulated mice. The data were shown as mean ± SD of triplicates experiments. *p < 0.05, **p < 0.01, *** p < 0.005 compared with the corresponding control. Differences in the results of two groups were evaluated using either two-tailed Student’s t-test or one-way ANOVA.

### LEF1 directly regulated the transcription of PD1

It is known that LEF1, homologous to TCF1, acts as a transcriptional factor to directly or indirectly regulate downstream genes in T cell development. To verify whether LEF1 directly regulates PD1 expression or not, we established a LEF1 knockdown (LEF1 KD) T cell line (MOLT4 cell line). In this cell line, LEF1 is highly expressed as well as TCF1. With the use of RNAi, LEF1 was downregulated by western blot (**Figure 5A**). By RT-qPCR, not only the mRNA level of LEF1 was downregulated, and also PD1 expression was reduced in LEF1 knockdown cells (**Figure 5B and 5C**). LEF1 positively regulated PD1 in human cell line, which was consistent with the results in the mouse models. It has been verified that LEF1 transcriptionally regulates several targeted genes, including CD4+ lineage-associated genes, such as St8sia6,Cd40lg and Itgb3, and CD8+ T cell effector genes Prdm1, Prf1 and Fasl [26]. LEF1 regulated these genes by recognizing and binding to a conserved sequence motif in the promoter regions of these genes, as it was shown (**Figure 5D**). We mutated the conserved potential LEF1-bound sequence “ATCAAAG” to “GCTGGGA” by converting “A” to “G” and “T” to “C” and inserted these promoter regions to luciferase reporter plasmids (**Figure 5E**). Luciferase activity of wildtype promoter of PD1 was higher than that of both mutated promoter of PD1 and control vector by the dual luciferase reporter assay (**Figure 5F**). It indicated that PD1 is likely to be transcriptionally regulated by LEF1. By performing ChIP-qPCR, we further determined that LEF1 directly regulated PD1 transcription (**Figure 5G**). According to the online database of ChIP-sequencing, we also confirmed that PD1 promoter strongly bound to LEF1 for transcription activation (**Figure 5H**). These results further corroborated the notion that LEF1 transcriptionally regulated PD1 by directly binding its promoter region of the genome.

**Figure 5.**
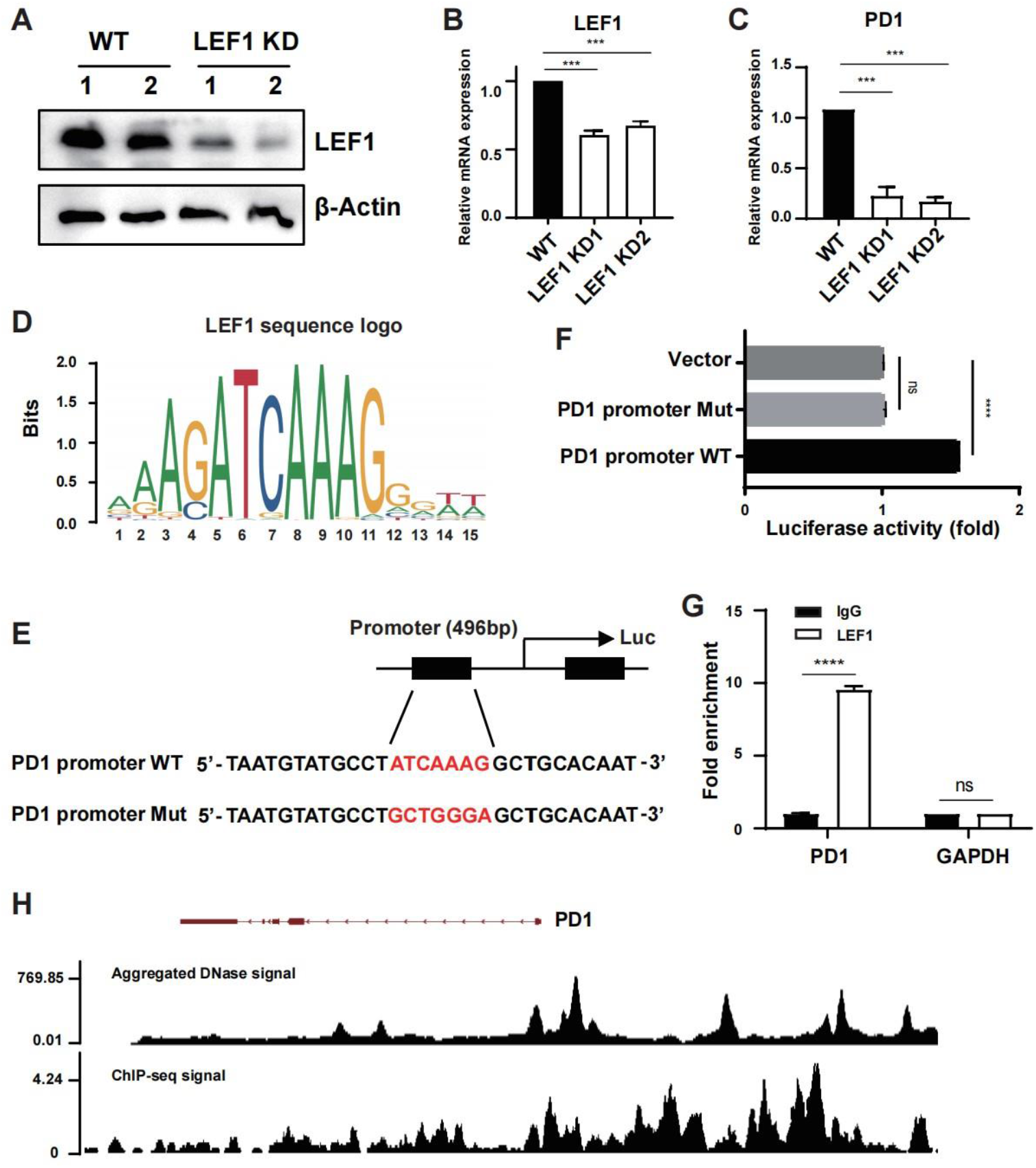
LEF1 transcriptionally regulated PD1. (A) Western blot of LEF1 in wild-type and RNAi MOLT4 cells. (B and C) RT-qPCR of LEF1 and PD1 in wild-type and LEF1 knockdown MOLT4 cells. (D) Sequence logo for conserved LEF1-binding sites. (E) The Determination of mutation point by sequence analysis in the promoter region of PD1. (F) Bar graph to examine the promoter activity of wild-type and mutated PD1 promoters. Vector alone in the reaction mixture was used as a negative control. (G) ChIP-qPCR of PD1 with the antibody of LEF1 and IgG control. (H) The online ChIP-seq data analysis of PD1. The online website was linked (https://www.factorbook.org/human/LEF1/mememotif/ENCSR240XWM/ENCFF420OHR/[ag]a[cg]atcaaagg/genomebrowser). The data were shown as mean ± SD of triplicates experiments. *p < 0.05, **p < 0.01, *** p < 0.005 compared with the corresponding control. Differences in the results of two groups were evaluated using either two-tailed Student’s t-test or one-way ANOVA.

## Discussion

In this work, we have presented the transcription regulation of PD1 by LEF1 in mature T cells, including CD8+ T cells and Treg cells, providing novel insight into the molecular biology of PD1.

The transcription factor LEF1, as well as TCF1, is vital for the development and functions of CD8+ T cells and Treg cells. TCF1 has been reported to transcriptionally repress the genes of which proteins are bound by FOXP3, although it does not alter the core Treg cell transcriptional signature [35]. TCF1 deficiency strongly inhibits T cell proliferation and cytotoxicity in Treg cells and promotes tumor growth. Besides TCF1, LEF1 knockout also leads to the absence of immune-suppressive capability of Treg cells. Both TCF1 and LEF1 do not perturb Treg cell homeostasis. High PD1 expression enhances the effector phenotype of Treg cells when the T cell receptor is activated [36]. Our results verify that LEF1 regulates PD1 in Treg cells and CD8+ cells. This is consistent with these previous studies to some extent. LEF1 increases PD1 expression and thus promotes the immune-suppressive capability of Treg cells. Further researches revealed that Treg cells can be divided into three different subsets, including the R1 subset (rTreg), R2 subset (aTreg), and R3 subset (eTreg), based on the gradient expression of TCF1 and LEF1 [37]. It is confirmed again that TCF1 and LEF1 are essential for the suppressive capacity of Treg cells to prevent autoimmunity. Among these subsets, only rTreg cells highly express LEF1. But there is no mention of PD1 expression in this subset. Although PD1 is dispensable for Treg cell development, PD1 can enhance or inhibit peripherally induced Treg cells [38,39]. It needs to be further explored.

Naive T cells express PD-1 rapidly when the TCR signaling pathway is activated. However, when T cells are continuously stimulated, they will develop into exhausted T cells and continuously express high PD-1. Besides CD4+ and CD8+ T cells, PD1 is also expressed in Treg, B, NK, and NKT cells. It is well known that T cell exhaustion limits immune responses against cancer and it is a major cause of resistance to chimeric antigen receptor [16]-T cell therapeutics. Further studies on the molecular characteristics associated with CAR T cell differentiation have revealed that after the CAR T cell infusion, the transition toward T cell exhaustion is blocked by the repression of genes (TCF7 and LEF1) with memory potential and DNA demethylation of genes (CX3CR1, BATF, and TOX) [40]. So, whether PD1 expression controlled by LEF1 in a particular condition is involved in this process is still unclear. Further studies have evidenced that there are two subgroups of exhausted T cells, namely PD1+ TCF1+ progenitor-like precursor exhausted cells and PD1+ TCF1-terminally differentiated exhausted T cells [41]. And the progenitor-like precursor cells still function in immunotherapy. However, it is interesting to know LEF1 levels in these two kinds of exhausted T cells. In addition, PD1 is highly expressed in exhausted CD8+ T cells, which explains why the ratio of PD1 in tumor CD8+ T cells is relatively higher than that of CD8+ T cells in the spleen on the whole.

Recent work has also shown that PD1 is strongly upregulated on T cells (CD4+ and CD8+ T cells) stimulation among bovine PBMCs with the treatment of ConA and becomes more in cattle with bovine leukemia virus infection [42]. ConA-stimulated porcine Cd8+ T cells highly express genes of TCF7 and LEF1 in vitro conditions, but it only occurs in the naïve CD8+ T -cell subsets [43]. Therefore, it is indicated that ConA stimulates the expression of both PD1 and LEF1 in some extent. With the treatment with ConA, the growth of the liver tumors is repressed in hepatoma-bearing SCID mice, but the highest dose of ConA is 20 mg/kg [44]. And when the dose of ConA is at 7.5 mg/kg, liver tumor formation is significantly reduced and mice survival is prolonged in hepatoma-bearing mice (ML-14a cells). It is regarded that ConA activates tumor-specific CD8+ CTLs and NK cells for T cell-mediated responses. In our study, ConA promotes PD1 expression in tumor-specific CD8+ and Treg CTLs, which may conflict with the results of this study to some extent.

In summary, our study has revealed the transcriptional regulation of TFC1 on PD1 in mature T cells, including CD8+ T cells and Treg cells. LEF1 regulated PD1 by directly binding to the conserved motif of its promoter regions. Because most researchers focus on PD1 expression in some specific conditions, it is a little difficult to uncover the transcription of PD1. Our findings highlight the pivotal regulation of a transcription factor, LEF1, in regulating PD1 expression to accelerate the progression of CD8+ T cell exhaustion and Treg cell effects.

## Declarations

### Ethics approval and consent to participate

All methods are reported in accordance with ARRIVE guidelines (https://arriveguidelines.org) for the reporting of animal experiments. The experimental procedures and animal care of this study were strictly carried out in accordance with the guidelines of the Animal Experiment Center of Guangdong Health Science.

### Consent for publication

Not applicable.

### Availability of data and materials

The data that support the findings from this study are available on request from the corresponding authors. The data are not publicly available due to privacy or ethical restrictions.

### Author contributions

P.Z. designed and performed the experiments, analyzed the data and prepared the figures. L.S. and C.Z. participated and interpreted the experiments. P.Z. wrote the draft. All authors critically revised the manuscript.

## Acknowledgements

We thank Yubei Jin and Hanlin Wang for the helps with Western blot. We thank Sangon Biotech (Shanghai) Co., Ltd for DNA synthesis and sequencing. We thank Dr. Zhigang Mai for the instructions of research-grade positive-mounted automatic fluorescence microscope at Shenzhen University Instrument Management Platform.

